# RAD: a web application to identify region associated differentially expressed genes

**DOI:** 10.1101/2020.08.03.234302

**Authors:** Yixin Guo, Ziwei Xue, Ruihong Yuan, William A. Pastor, Wanlu Liu

## Abstract

With the advance of genomic sequencing techniques, chromatin accessible regions, transcription factor binding sites and epigenetic modifications can be identified at genome-wide scale. Conventional analyses focus on the gene regulation at proximal regions; however, distal regions are usually neglected, largely due to the lack of reliable tools to link the distal regions to coding genes. In this study, we introduce RAD (Region Associated Differentially expressed genes), a user-friendly web tool to identify both proximal and distal region associated differentially expressed genes. RAD maps the up- and down-regulated genes associated with any genomic regions of interest (gROI) and helps researchers to infer the regulatory function of these regions based on the distance of gROI to differentially expressed genes. RAD includes visualization of the results and statistical inference for significance.

**Availability:** RAD is implemented with Python 3.7 and run on a Nginx server. RAD is freely available at http://labw.org/rad as online web service.

## Introduction

Data-rich methods such as micrococcal nuclease sequencing (MNase-seq, Schones, et al., 2008), DNase I sequencing (DNase-seq, Boyle, et al., 2008), chromatin immunoprecipitation sequencing (ChIP-seq, Barski, et al., 2007) assay for transposase-accessible chromatin sequencing (ATAC-seq, Buenrostro, et al., 2013) and whole genome bisulfite sequencing (WGBS, Cokus, et al., 2008) that analyze genome-wide epigenetic landscape have been widely used to provide information on the binding of transcription factors (TFs) and chromatin accessibility of cis-regulatory elements (CREs) including promoters and enhancers. Through peak or DMR (differential methylated regions) calling, one can identify genomic regions of interest (gROI) in genomic data, which provides the basis for further analysis. Integration of these methods with RNA sequencing (RNA-seq) data allows researchers to determine whether differentially expressed genes (DEGs) are regulated by TF binding, chromatin accessibility, or other epigenetic modifications such as DNA methylation.

Methods for the integrative analysis of multi-level omics data have been introduced in recent years, such as GREAT, which incorporates ChIP-seq data and gene ontologies to highlight the association between CREs and gene function (McLean, et al., 2010). BETA, a new generation tool incorporates transcriptome and ChIP-seq data to infer direct target genes (Wang, et al., 2013). Combination of multi-level omics data may provide new insights into the regulation of transcription by genome-wide epigenetic landscape.

Conventional analyses that focus on proximal regulatory events often omit information distal to gROI. Unlike promoters that are adjacent to transcription start site (TSS) (≤ 1kb), enhancers may activate their target promoters and regulate the expression of target genes from long distance (Shlyueva, et al., 2014). In this study, we introduce a user-friendly web application, Region Associated DEGs (RAD), to intuitively measure both proximal and distal association between TF binding, chromatin accessibility, epigenetic modification or any other gROI and the transcriptional changes of surrounding genes. Using a hypergeometric test, we can potentially infer whether nearby genes are up-regulated or down-regulated by differential TF binding, chromatin accessibility or epigenetics changes, and whether this regulation is mediated via proximal and/or distal interaction. The algorithm used in RAD has been successfully implemented in recent publications to investigate the association of DEGs with chromatin accessibility changes (Pastor, et al., 2018), TF binding (Harris, et al., 2018) and DMRs (Gallego-Bartolome, et al., 2019). The web application thus allows users to infer potential regulatory effects of transcription factors, epigenetic modifications or any gROI in genomic data.

## RAD functions

RAD is an open-access, user-friendly web application (Supplemental Figure 1-4) for studying the relationship between gROI and DEGs. Visualization of DEGs surrounding gROI are implemented in RAD to help researchers to infer the potential regulatory function of transcription factors or chromatin features.

## RAD Input

The input files for RAD include three files: 1-2) line-break text file containing up- or down-DEGs (*upregulated_genes.txt, downregulated_genes.txt*); 3) file containing gROI information in browser extensible data format (*gROI_file.bed*). Up- or down-regulated genes can be calculated with DEseq2 (Love, et al., 2014) or other methods that identify differentially expressed genes. Genes in the up- or down-DEGs files should be separated by line breaks (i.e. each line should only contain one gene symbol or Ensembl ID). gROI file provides the genomic region information including chromosome number, start and end position of the region. Instead of uploading the data, the user can also directly paste a list of gene names and genomic regions into the text-input area on the website.

Required options include user defined reference genome and gROI extended distance. Reference genome corresponding to the data should be specified by user and we support several widely used mammalian (*Homo sapiens, Mus musculus*) and plant (*Arabidopsis thaliana*) reference genomes, including GRCh38, GRCh37, GRCm38, GRCm37 and TAIR10 (www.ensembl.org). gROI extended distance can be chosen from 1kb, 10kb, 25kb, 50kb, 100kb, 500kb and 1000kb with 1000kb as default. The title and color palette of the output bar plot can be customized according to user’s preference.

## RAD Workflow and implementation

RAD web application was implemented with Python (version 3.7) programming language, on a Nginx server with Centos 7.06 operating system. The website was developed using AngularJS and Flask framework. The algorithm can be divided into four steps (Figure 1). The first step is to identify gROI associated genes within user defined gROI extended distance through *awk* and *bedtools* (Quinlan and Hall, 2010). Then gROI extended distance will be split into different distance bins. Up- or down-regulated genes are then mapped into different distance bins. To calculate the enrichment of up- or down-regulated genes within different distance bins, observed over expected ratio is calculated as indicated in Figure 1. Genes that are outside of the gROI extended distance (too far from gROI), or not differentially regulated are excluded. The third step is to perform hypergeometric test to calculate the *p-value*.

**Figure 1.**
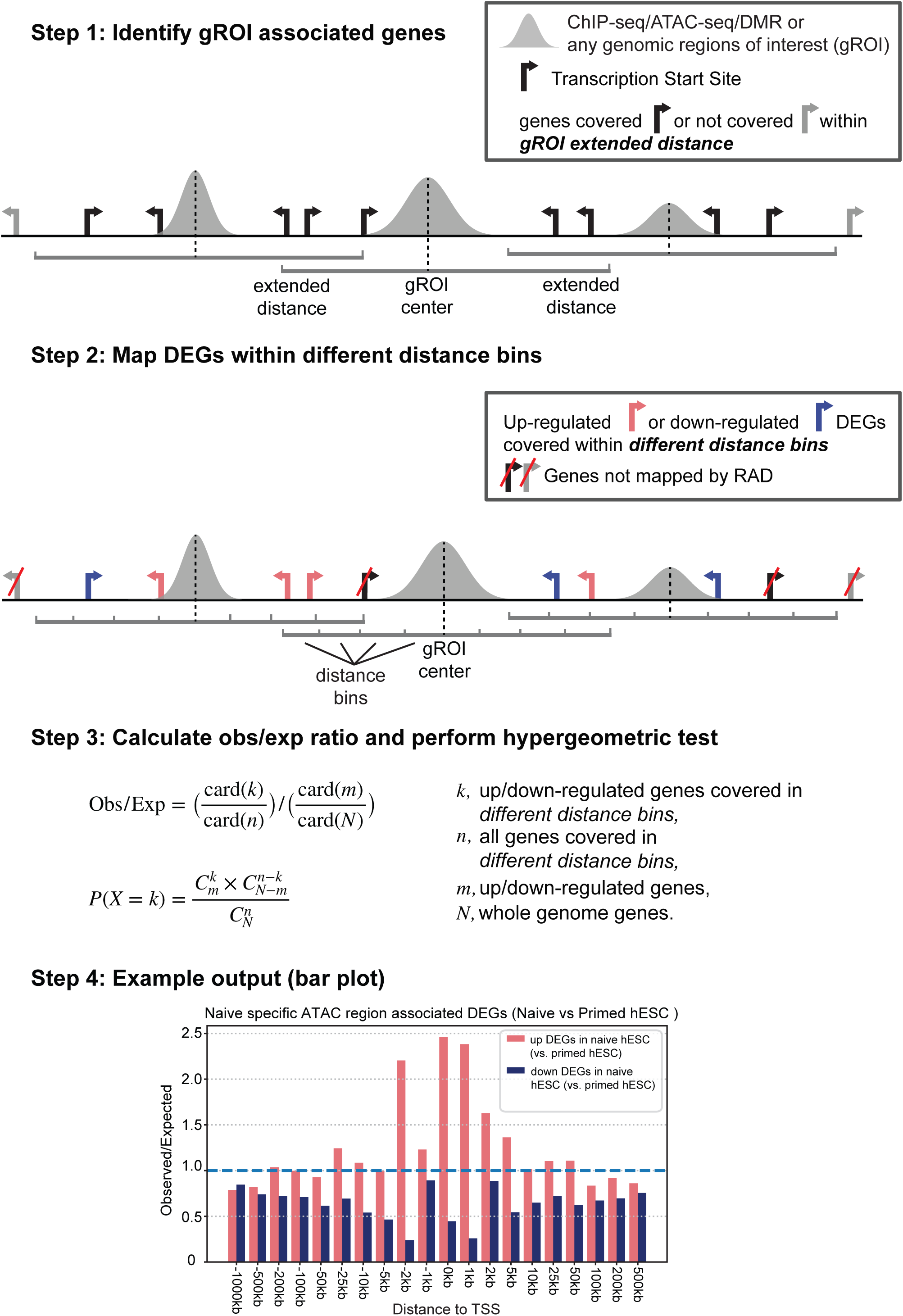
Major steps of the algorithm implemented in RAD. The algorithm includes Step 1) identify of gROI associated genes; Step 2) map DEGs within different distance bins; Step 3) calculate observed over expected ratio and perform hypergeometric test; Step 4) example output bar plot (Data from *Pastor, et al., 2018*).

Finally, DEGs covered by gROI extended distance, count of up- or down-regulated DEGs in each distance bins and the calculated *p-value* will be reported in text files. The observed over expected ratio will be displayed on the website as bar plot. An example output bar plot comparing naïve human embryonic stem cells (hESC) specific ATAC-seq peaks and naïve hESC up- or down-DEGs is displayed in Figure 1, suggesting the potential proximal and distal transcriptional promotion role of those naïve specific ATAC-seq peak (Data from Pastor, et al., 2018).

## RAD Output

RAD output contains three files: 1) DEGs covered by extended gROI are stored in a text file named *RAD_genename_distance.txt*; 2) The count of up- or down-regulated DEGs in each distance bins, total genes count genome-wide as well as the calculated *p-value* will be reported in a text file named *RAD_genecount_pvalue.txt*; 3) The bar plot of observed over expected ratio in each distance bins can be downloaded as png, SVG or pdf format.

## Conclusion

We developed a web application RAD to identify gROI associated DEGs and provide a graphic output as well as gROI associated DEGs list for further analysis. Downstream analysis such as gene ontology (GO) enrichment analysis for DEGs in certain distance bins could be performed to help biologists to further infer potential functions of gROI.

## Acknowledgements

We thank Dr. Chen, Di, Dr. Wang, Chaochen from Zhejiang University and Dr. Li, J. Jessica from UCLA for helpful discussion. This work is supported by Zhejiang Provincial Natural Science Foundation of China, [LQ20C060004 to W.L.]; and the Fundamental Research Funds for the Central Universities[K20200099]. W.L., W.P. developed the algorithm. Y.G., Z.X. developed the web application. R.Y. implemented the algorithm in python. Y.G., Z.X., W.L. wrote manuscript and W.L. coordinated research.

