## Supplementary Figures for "RAD: a web application to identify region associated differentially expressed genes"

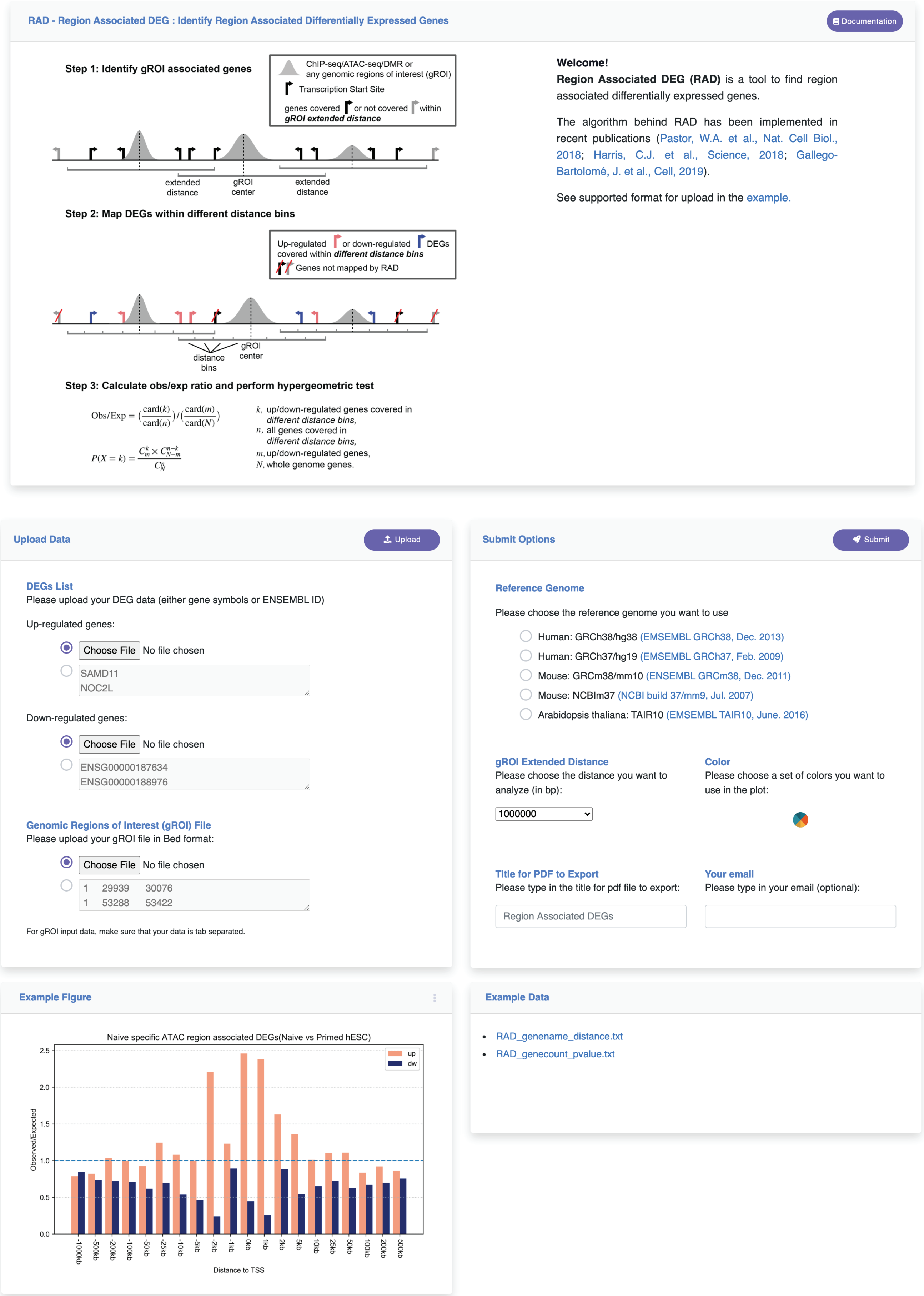


**Supplemental Figure 1. Overview of the web user interface for RAD.** RAD is freely available at *http://labw.org/rad* as online web service.

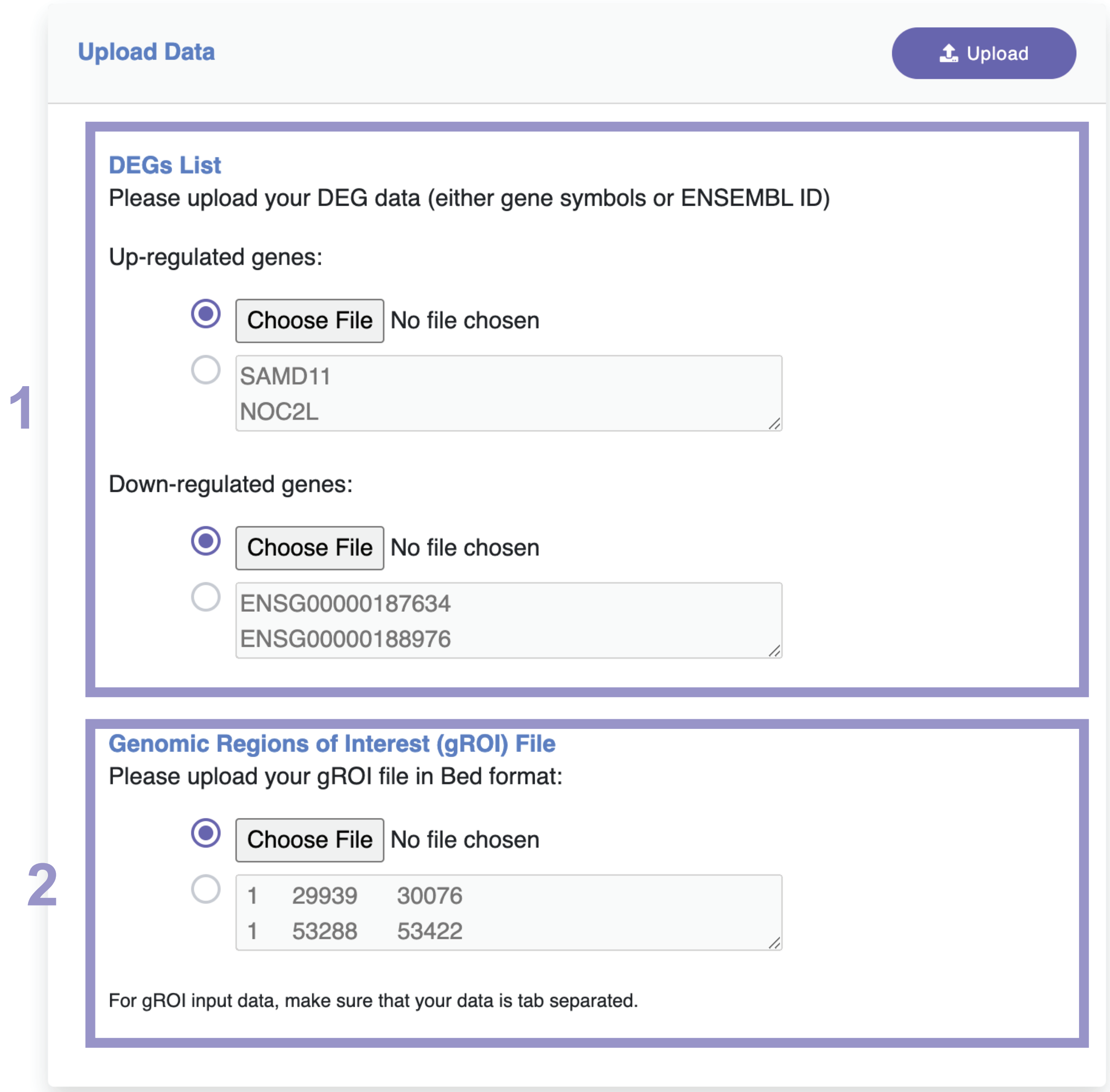


**Supplemental Figure 2. The file-submission component of RAD.** The file submission interface has two sections: 1. Uploading differentially expressed genes list in the format of *.txt* or *.csv* files; 2. uploading gROI file in the format of .bed file. Another option is to directly paste a list of gene names (either gene symbol or Ensembl ID) and genomic regions into the text-input area on the website.

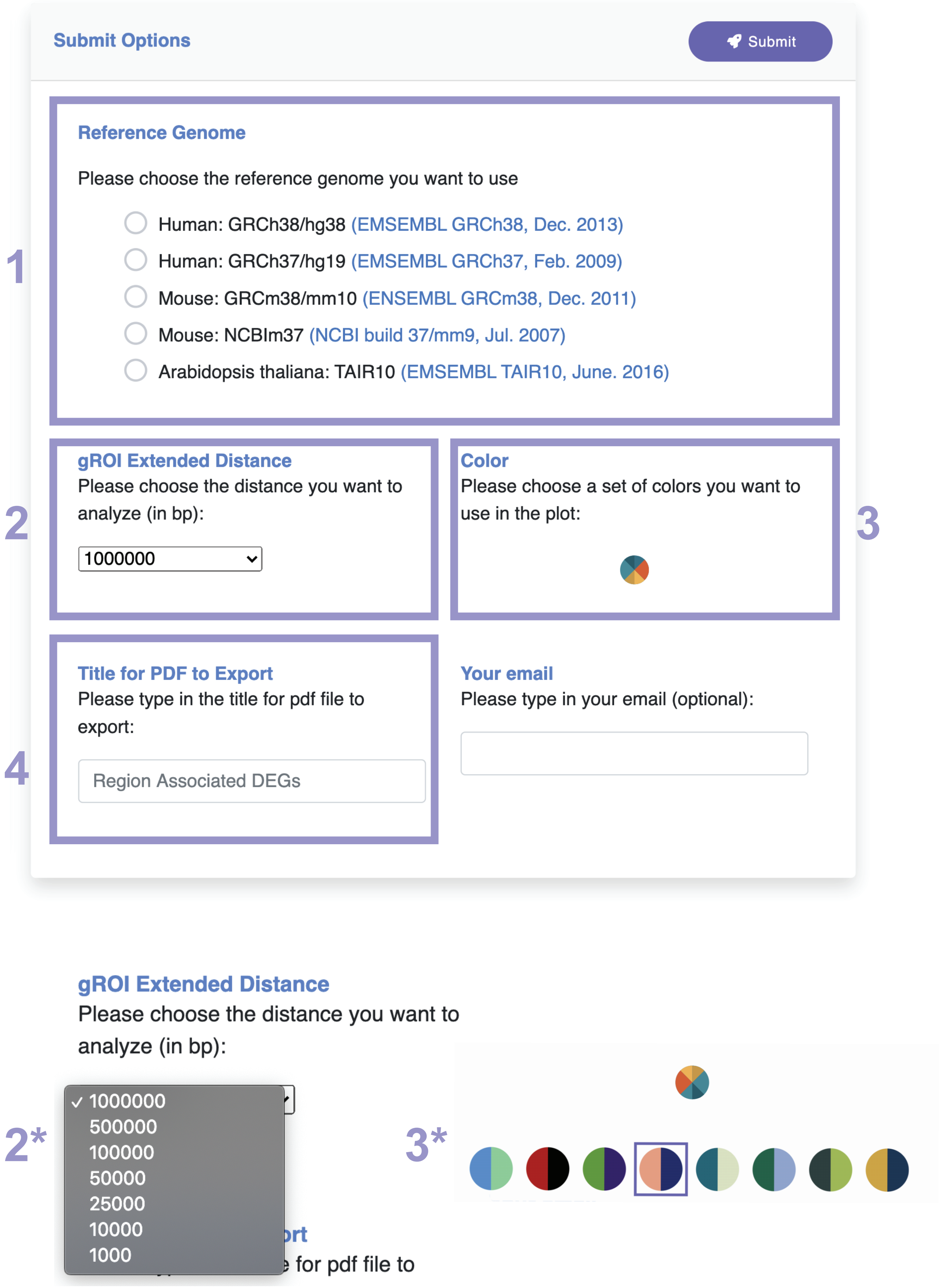


**Supplemental Figure 3. The submission options component of RAD.** There are four sections for submission options: 1. specify the reference genome (required); 2. choose the gROI extended distance (2*, drop-down list of seven options are provided with 1000kb as default); 3. pick a set of colors to be used in the plot (3*, pop-up list of eight color palette are provided with pink and purple color palette as default); 4. enter the title for bar plot.

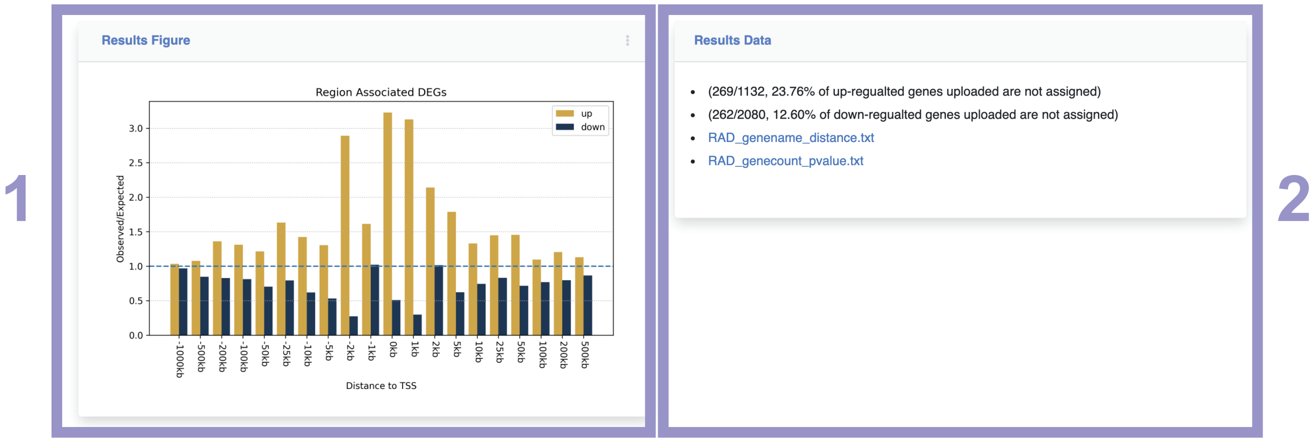


**Supplemental Figure 4. The output component of RAD.** There are two sections for the output: 1. The observed over expected ratio bar plot in different distance bins; 2. the output text files containing DEGs covered by extended gROI (*RAD_genename_distance.txt*) and count of up- or down-regulated DEGs in each distance bins, total genes count genome-wide, as well as the calculated p-value (*RAD_genecount_pvalue.txt*).
